# Expression of *SULTR2;2* in the Arabidopsis bundle sheath is mediated by a highly conserved positive regulator

**DOI:** 10.1101/250738

**Authors:** Sandra Kirschner, Helen Woodfield, Katharina Prusko, Maria Koczor, Udo Gowik, Julian M. Hibberd, Peter Westhoff

## Abstract

The bundle sheath provides a conduit linking veins and mesophyll cells. In C_3_ *Arabidopsis thaliana* it also plays important roles in oxidative stress and sulphur metabolism. However, the mechanisms responsible for the patterns of gene expression that underpin these metabolic specialisations are poorly understood. Here we used the *A. thaliana SULTR2;2* gene as a model to better understand mechanisms that restrict expression to the bundle sheath. Deletion analysis indicated that the *SULTR2;2* promoter contains a short region necessary for expression in the bundle sheath. This sequence acts as a positive regulator and is tolerant to multiple consecutive deletions indicating considerable redundancy in the *cis*-elements involved. It is highly conserved in *SULTR2;2* genes of the Brassicaceae and is functional in the distantly related C_4_ species *Flaveria bidentis* that belongs to the Asteraceae. We conclude that expression of *SULTR2;2* in the bundle sheath is underpinned by a highly redundant sequence that likely represents an ancient and conserved mechanism found in families as diverse as the Asteraceae and Brassicaceae.

## Introduction

The evolution of multicellularity is associated with individual cell-types being able to undertake specialised roles within a tissue. In leaves, bundle sheath (BS) cells form a wreath-like structure around the vasculature that appears analogous to the endodermis of roots (Esau, 1965). The role of BS cells is best characterized in C_4_ species that partition photosynthesis between the BS and mesophyll cells. In most C_4_ plants, after HCO_3_ - is initially fixed into C_4_ acids by phospho*enol*pyruvate carboxylase in mesophyll cells, these C_4_ acids then diffuse to the BS where CO_2_ is released and refixed by Ribulose-1,5-Bisphosphate Carboxylase/Oxygenase (RuBisCO). Decarboxylation of C_4_ acids in the BS generates a high concentration of CO_2_ around RuBisCO that suppresses the oxygenase activity of the enzyme and in so doing reduces photorespiration (Hatch, 1987). Thus, in C_4_ species, the BS is specialized to allow efficient fixation of CO_2_ in the Calvin-Benson-Bassham cycle. In some C_4_ plants, the BS is also modified in terms of light capture. For example, in maize and sorghum Photosystem II does not fully assemble in the BS (Kubicki *et al*., 1994) but components of cyclic electron transport are more abundant in the BS compared with mesophyll cells (Takabayashi *et al*., 2005). In addition to these changes associated with photosynthesis, the C_4_ BS is also modified to preferentially undertake starch synthesis and degradation, as well as the initial steps of sulphur assimilation (Friso *et al*., 2004).

In C_3_ plants, the role of the BS is less clearly defined. It is thought to help maintain hydraulic integrity of the xylem (Sade *et al*., 2014), regulate flux of metabolites in and out of the leaf (Leegood, 2008) and act as a starch store (Miyake and Maeda, 1976). The C_3_ BS is less important for photosynthesis than that of C_4_ species. However, although only around 15% of all chloroplasts of the C_3_ leaf are found in bundle sheath cells (Kinsman and Pyke, 1998), reducing photosynthesis in these cells compromises growth and seed production (Janacek *et al*., 2009). Thus, although less photosynthetic than the C_4_ BS, this physiological analysis indicates that the BS of C_3_ plants is also specialized. This notion is consistent with analysis of gene expression in this cell type. For example quantification of transcripts available for translation indicate that the *A. thaliana* BS is likely important in sulphur metabolism, glucosinolate biosynthesis, trehalose metabolism and detoxification of active oxygen species (Aubry *et al*., 2014). In summary, in both C_3_ and C_4_ plants mechanisms must operate to restrict the expression of some genes to BS cells.

To date, most studies of the mechanisms responsible for preferential gene expression in the BS have occurred in C_4_ species (Hibberd and Covshoff, 2010). In the C_4_ dicotyledon *Flaveria trinervia* the glycine decarboxylase P-subunit (*GLDPA*) gene contains two promoters, one proximal to the coding region, and the other more distal. Activity of the distal promoter is high but not cell-type specific. However, in the presence of the proximal promoter, transcripts derived from the distal promoter are degraded in mesophyll cells by nonsense-mediated RNA decay of incompletely spliced transcripts (Engelmann *et al*., 2008; Wiludda *et al*., 2012). Despite the phylogenetic distance between the Brassicaceae and the Asteraceae the *GLDPA* promoter from *F. trinervia* is able to generate BS-specific activity in C_3_ *A. thaliana* (Engelmann *et al*., 2008; Wiludda *et al*., 2012). In *Amaranthus hybridus* 5’ and 3’ untranslated regions of the *RbcS1* gene act to restrict accumulation of the glucoronidase (GUS) reporter to the BS of C_4_ *F. bidentis* and appear to function as enhancers of translation (Patel *et al*., 2006). Lastly, in C_4_ *Gynandropsis gynandra* preferential expression of *NAD-ME1&2* genes in the BS is associated with coding sequence rather than UTRs or promoter elements (Brown *et al*., 2011). The motifs underpinning this regulation are a pair of duons that play a dual role in coding for amino acids as well as the spatial patterning of gene expression associated with the C_4_ leaf. Although these duons are present in C_3_ *A. thaliana* and many other land plants they do not act to generate cell-specific expression in the ancestral C_3_ state (Brown *et al*., 2011; Reyna-Llorens *et al*., 2016). In summary, current evidence indicates that gene expression in the BS of C_4_ species is controlled by a variety of mechanisms, some of which involve regulatory codes that are derived from the ancestral C_3_ state.

However, our understanding of how gene expression is restricted to the BS in C_3_ species is poor. A small number of promoters including *SHORT-ROOT* (Dhondt *et al*., 2010), *SCARECROW* (Wysocka-Diller *et al*., 2000) and *SULTR2;2* (Takahashi *et al*., 2000) have been reported to drive BS specific expression in *A. thaliana,* but the molecular nature of *cis*-regulatory elements controlling their expression is unclear. An increased understanding of these processes would not only advance our understanding of mechanisms underpinning cell specific gene expression in multicellular leaves, but also provide insight into whether C_4_ gene expression is built on pre-existing mechanisms found in C_3_ species.

To better understand mechanisms associated with gene expression in BS cells of C_3_ *A. thaliana* we analysed the *SULTR2;2* gene that encodes a low-affinity sulphur transporter (Takahashi *et al*., 2000). Elements in the promoter sequence that regulate the spatial patterning, and the strength of gene expression were identified. Specifically, preferential expression in the BS is mediated by a repetitive region that is highly conserved within orthologous genes from species of the Brassicaceae. This region acts to enhance expression in the BS rather than repressing expression in mesophyll cells. Furthermore, the *SULTR2;2* promoter from *A. thaliana* generates BS specificity in the C_4_ species *F. bidentis* that belongs to the Asteraceae. The most parsimonious explanation for this finding is that a common transcription factor is shared by these phylogenetically dispersed species, and that it functions in both the C_3_ and C_4_ BS.

## Results

Nucleotides −2815 to +123 relative to the predicted *SULTR2;2* translational start site have previously been reported to generate expression in the BS of *A. thaliana* (Takahashi *et al*., 2000). We confirmed this finding (Figure 1A-C). To test if sequence after the predicted start codon is required for expression in the BS a construct that terminated at nucleotide −1 relative to the predicted translational start was generated (Figure 1D). Staining showed that each construct led to strong accumulation of GUS in the BS of mature rosette leaves (Figure 1B,C,E,F). Staining was evident in vascular tissue as well as the BS but there was no evidence that GUS accumulated in mesophyll or epidermal cells from either construct (Figure 1B,C,E,F). Promoter activity was quantified using the MUG fluorimetric assay. GUS activity driven by the construct that contained 603 fewer base pairs at the 5’ end but an additional 123 nucleotides of coding sequence was about tenfold higher (Figure 1G). This difference could either be due to a negative regulator located between −3418 and −2185 nucleotides, or a positive regulator in sequence downstream of the predicted translational start site. A translational fusion between the yellow fluorescent protein (YFP) and the nuclear localized Histone 2B protein under control of the *SULTR2;2* promoter (Figure 1H) labelled nuclei of BS cells and vascular tissue (Figure 1I) and indicated that the presence of GUS in the BS was due to gene expression and not diffusion of the dye outwards from to vascular tissue. Consistent with previous reports (Chytilova *et al*., 1999) it was noticeable that vascular nuclei were elongated and rod-like compared with the more spherical ones in the BS (Figure 1I). We conclude that elements between nucleotides −2815 and the translational start site of *SULTR2;2* are sufficient to drive gene expression in vascular and BS cells of *A. thaliana*.

**Figure 1:**
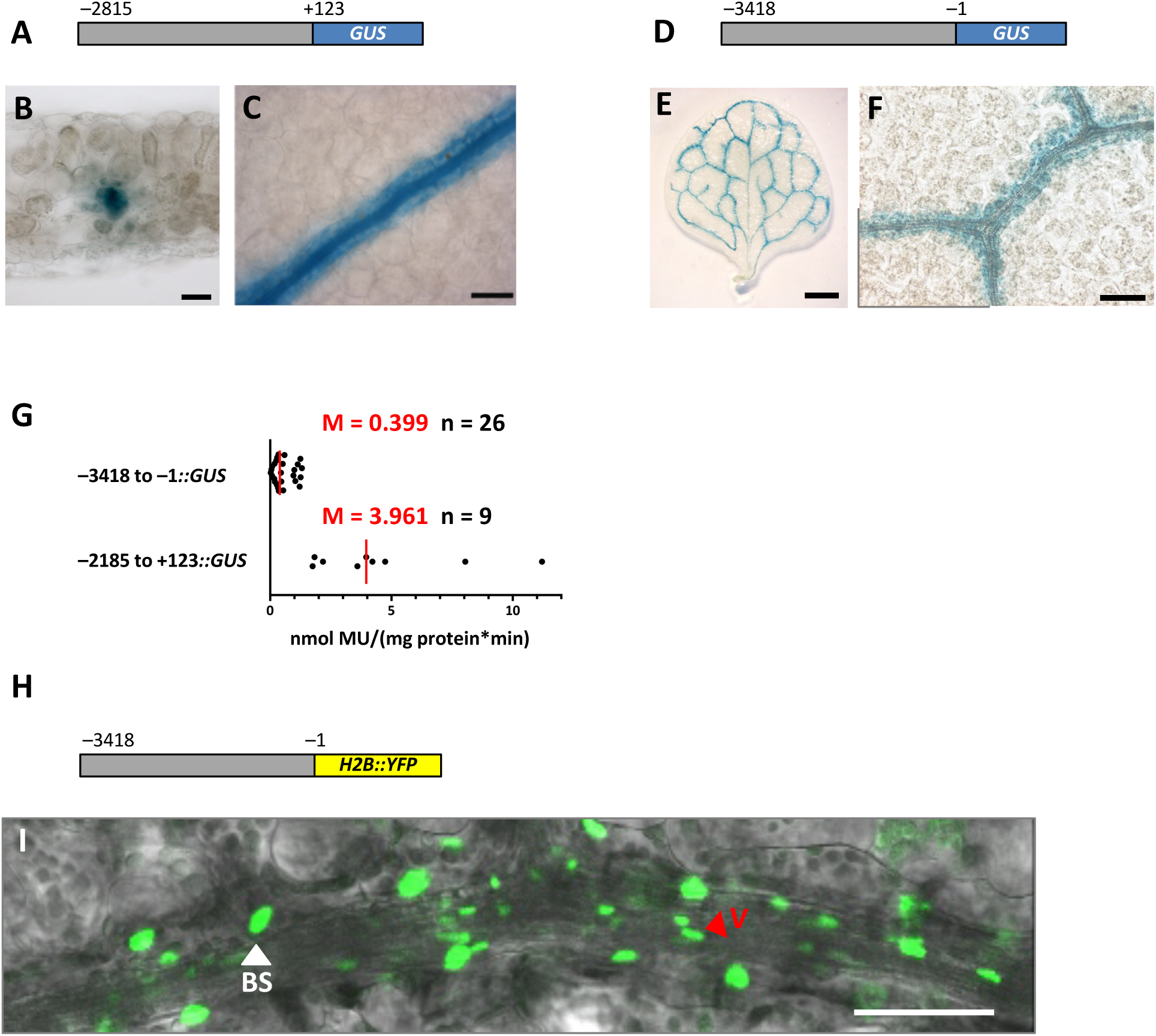
The *SULTR2;2* promoter drives expression in veins and bundle sheath cells of *Arabidopsis thaliana.* **(A-C)** Nucleotides −2185 to +123 of *SULTR2;2* are sufficient to generate preferential accumulation of the GUS reporter in bundle sheath cells. **(D-F)** Nucleotides −2185 to +123 of *SULTR2;2* are sufficient to generate preferential accumulation of the GUS reporter in bundle sheath cells. **(G)** Quantitative analysis of expression from each construct based on the GUS activity assay. **(H)**. Schematic of the *A. thaliana SULTR2;2* full-length promoter fused to *H2B::YFP* that targets YFP to the nucleus. **(I)** H2B::YFP fusion marks nuclei from the larger bundle sheath exhibits (white arrowhead) as well as the smaller more elongated nuclei of the vasculature (red arrowhead). Data from GUS activity assays include the median (M) indicated by red lines and the number (n) of independent lines. Histological GUS assays were allowed to proceed for 23 h (B), 16 h (C) and 5 h (E) 6 h (F). Scale bars represent 500 (C) or 50 µm (D,E,F).

### A short region that is necessary and sufficient to activate gene expression in the BS

To better understand the sequences responsible for generating expression in BS cells, a 5’ deletion series was generated (Figure 2A). Each of the deleted regions are hereafter referred to as regions 1 to 5. Removal of region 1 had no significant effect on either activity or spatial accumulation of GUS (Figure 2A&C) indicating that no essential *cis*-regulatory elements are located within this section. Deletion of region 2 resulted in total loss of GUS activity and staining (Figure 2A&D), and removal of regions 3 and 4 had no further effect (Figure 2A,E&F). These data therefore indicate that nucleotides in region 1 do not impact on promoter activity, but that region 2 is necessary for BS expression in mature leaves. A separate deletion series involving slightly smaller regions supported this conclusion, with expression in the BS being lost once nucleotides between −1813 and −1335 were deleted (Supplementary Figure 1).

**Figure 2:**
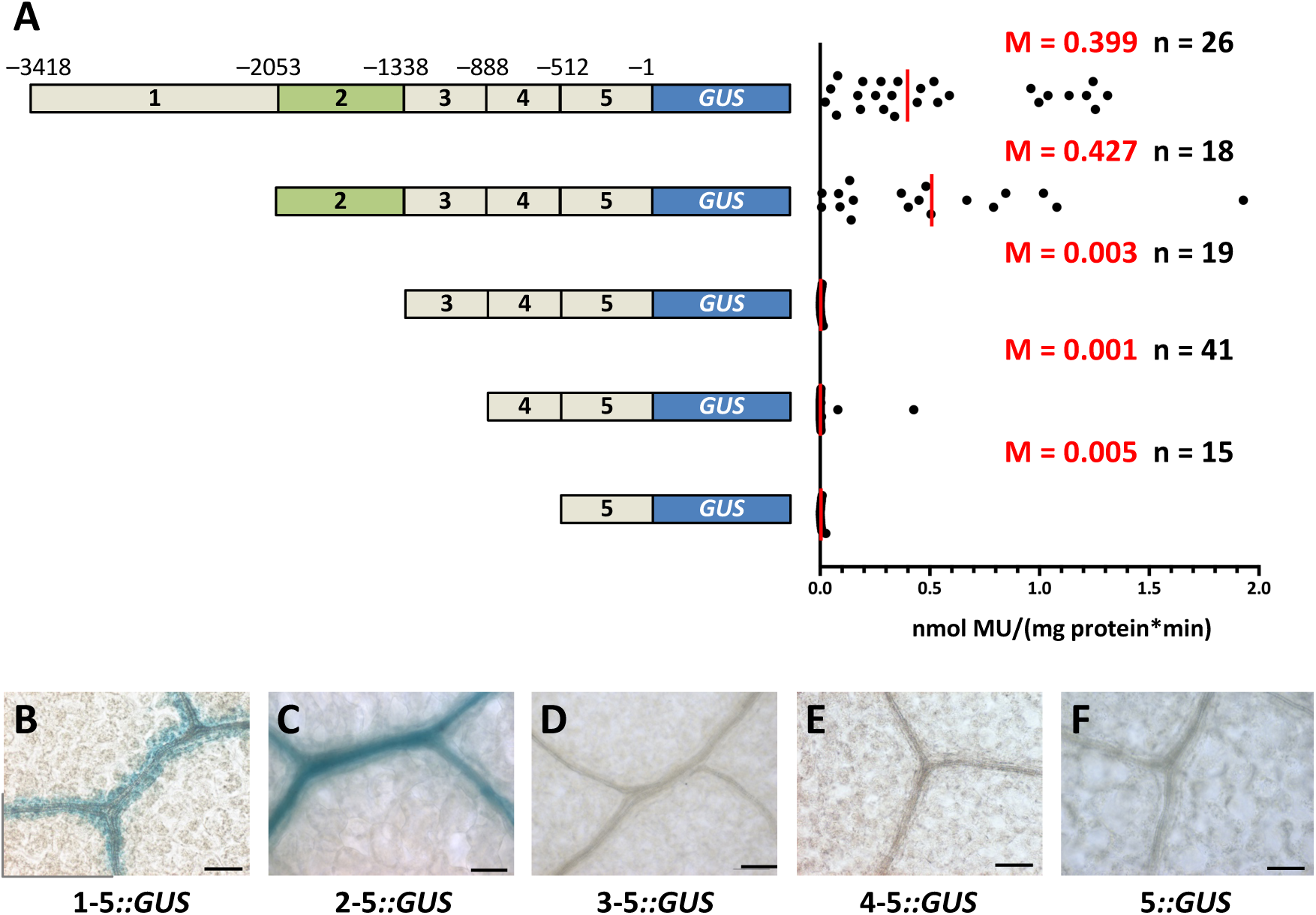
A 715 nucleotide region in the *SULTR2;2* promoter that is necessary for bundle sheath expression. **(A)** Schematic of the 5’ deletion series (left) with GUS activities (right). **(B-F)** GUS accumulation from each deletion. Deletion of region 2 abolished accumulation of GUS. Data from GUS activity assays include the median (M) indicated by red lines and the number (n) of independent lines. Histological GUS assays were conducted on leaves for 23 h (B), 4 h (C) and 6 days. Histological GUS assays were allowed to proceed for 23 h (B), 4 h (C) and 6 days (E&F). Scale bars represent 50 µm.

The lack GUS accumulation in the BS and loss of promoter activity once region 2 is removed could be because this region contains *cis*-elements that generate expression specifically in BS cells or because it drives ubiquitous expression but regions 3 to 5 contain elements that restrict activity to the BS. To investigate this possibility, 5’ rapid amplification of cDNA ends was first used to define the transcriptional start site of *SULTR2;2* (Supplemental Figure 2). No single strong transcription start site was detected but rather multiple transcripts were initiated from position −125 onwards (Supplementary Figure 2). Nucleotides spanning −349 to −1 were therefore considered likely to be sufficient for transcriptional initiation and are hereafter referred to as the core promoter. Fusion of region 2 to this core promoter led to MUG conversion that was comparable to that from the full-length promoter (Figure 3A and Figure 2A) and was also sufficient to direct accumulation of GUS to veins and the BS (Figure 3B). To exclude the possibility that the core promoter includes *cis*-elements necessary for BS-specific expression, region 2 was also fused to the minimal CaMV35S promoter which does not drive significant expression. Although this led to fourfold lower GUS activity than the full-length promoter (Figure 3A) GUS accumulation was still restricted to veins and the BS (Figure 3C). We conclude that although nucleotides −349 to −1 are not able to generate expression in the BS (Figure 2F), they likely represent the core promoter of *SULTR2;2*. In contrast, nucleotides −2053 to −1312 are sufficient to restrict expression to the BS and vascular tissue of *A. thaliana*.

**Figure 3:**
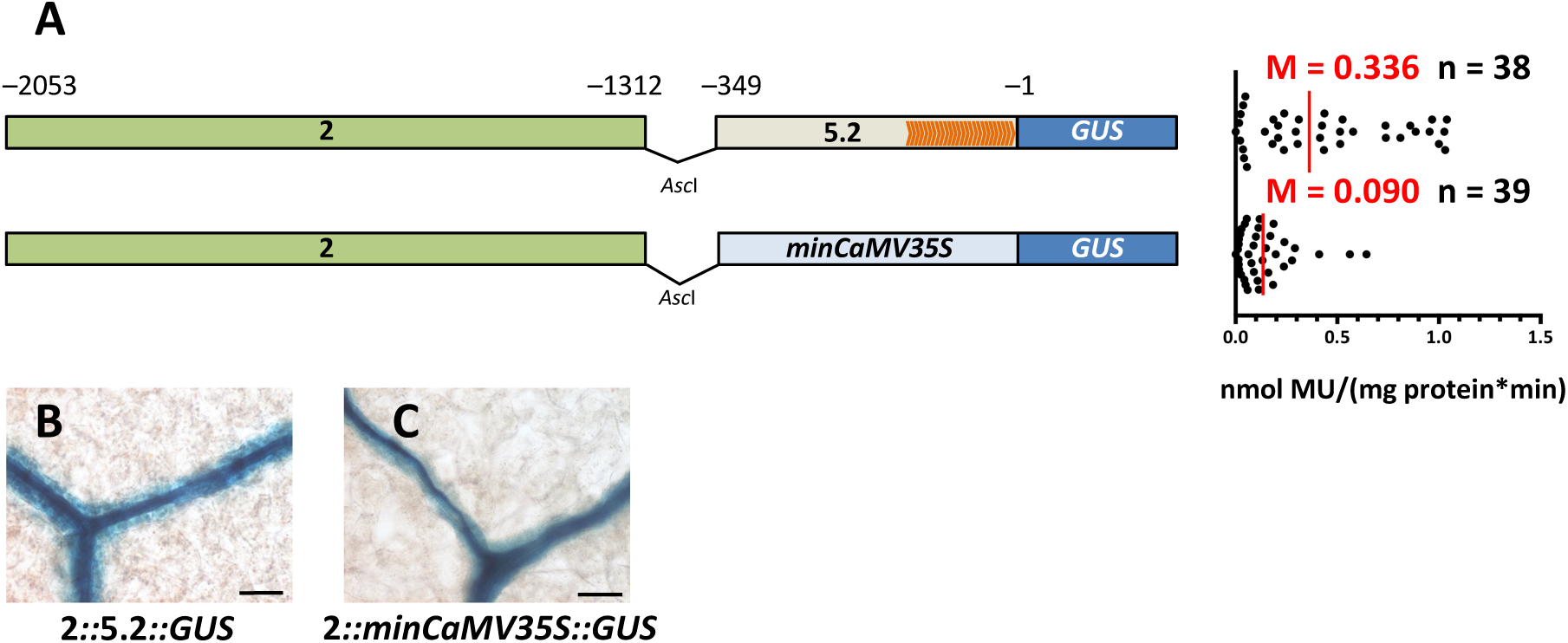
The *SULTR2;2* promoter contains a region that is sufficient to activate expression in bundle sheath cells. **(A)** Schematic of constructs containing nucleotides −2053 to −1312 combined with either the core promoter of *SULTR2;2* or the minimal CaMV35S promoter (left) and quantitative analysis of expression from each construct based on the GUS activity assay (right). Orange arrowheads within the core promoter indicate transcription start sites obtained by 5’-RACE. **(B-C)** Both constructs are sufficient to generate GUS accumulation in the bundle sheath. Data from GUS activity assays include the median (M) indicated by red lines and the number (n) of independent lines. Histological GUS assays were allowed to proceed for 5 h (B) and 22 h (C). Scale bars represent 50 µm.

### *AtSULTR2;2* contains multiple redundant sequences mediating BS expression

Having established that nucleotides −2053 to −1312 relative to the predicted translational start site are both necessary and sufficient for BS expression, an unbiased approach to further dissect this region was adopted. Ten consecutive deletions were made to this region and each fused to the core promoter of *SULTR2;2* (Figure 4A). Strikingly, none of these deletions resulted in total loss of GUS activity (Figure 4A), nor was GUS staining lost from the BS (Figure 4B-K) indicating that despite an absence of repeated *cis-*elements in this region significant functional redundancy in regulatory elements must mediate BS-specific expression. However, it was notable that GUS activity declined to varying degrees compared with the intact region (Figure 4A), suggesting that either these redundant *cis*-elements act additively, or that this region contains quantitative elements regulating gene expression.

**Figure 4:**
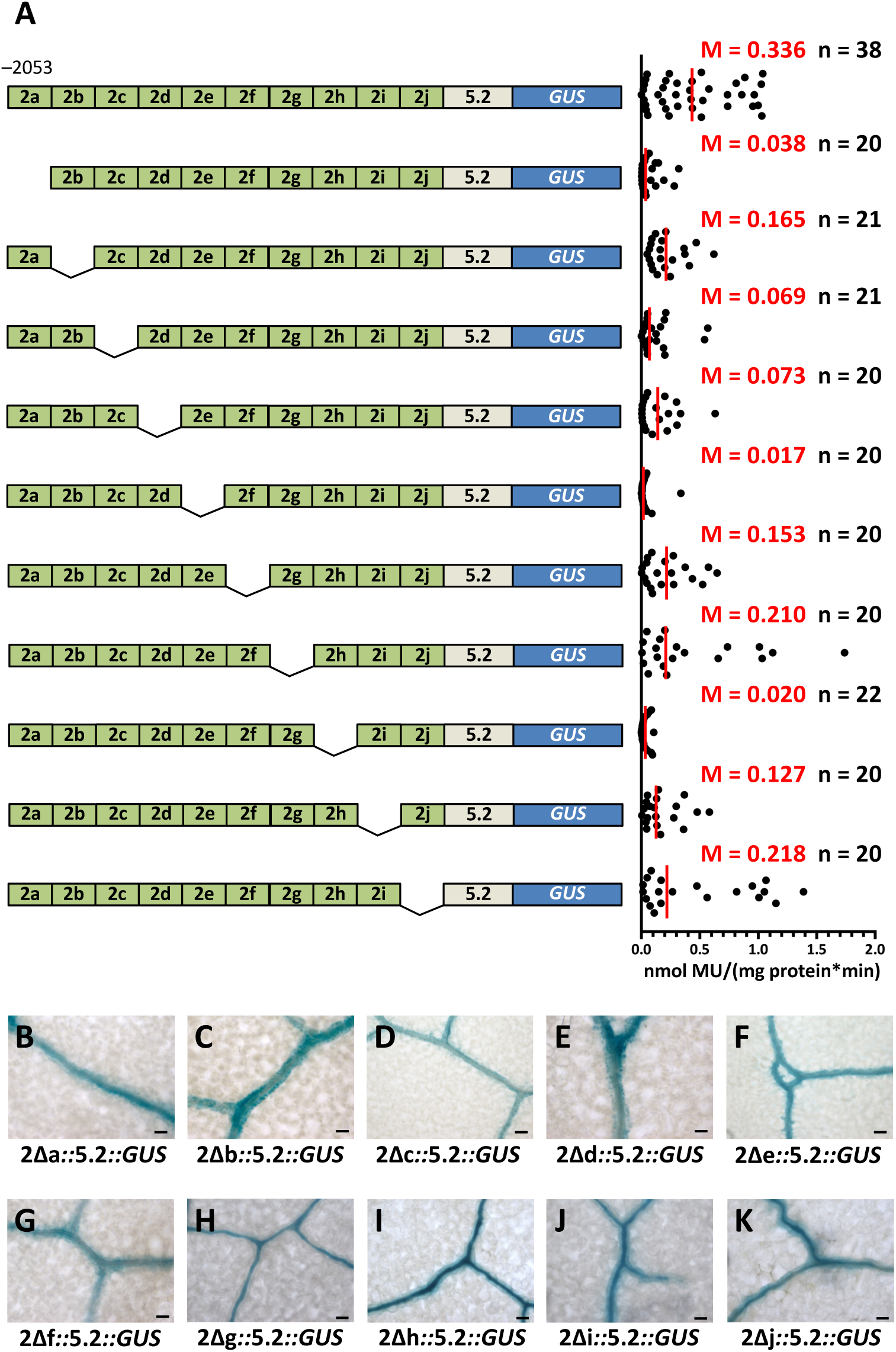
*SULTR2;2* contains a region with multiple redundant regions capable of directing bundle sheath expression. **(A)** Schematic of internal deletion series constructs (left) and quantitative analysis of expression from each construct based on the GUS activity assay (right). Except for the last deletion representing 66 base pairs, each construct lacks consecutive 75 base pair sequences. These internal deletions modify the GUS activity **(A)**, but none abolish accumulation of GUS in the bundle sheath **(B-K)**. Data from GUS activity assays include the median (M) indicated by red lines and the number (n) of independent lines. Histological GUS assays were allowed to proceed for 23 h (B), 16 h (C), 48 h (D), 23 h (E), 48 h (F), 8 h (G), 3 h (H), 7 h (I), 3 h (J) and 19 h (K). Scale bars represent 50 µm.

To better understand the extent to which *cis-*elements in this section of the promoter act redundantly, larger deletions were made from position −2053 (Figure 5A). This generated five regions, hereafter referred to as sub-regions 2.1 to 2.5. Deletion of sub-region 2.1 resulted in a strong reduction of GUS activity (Figure 5A) but BS-specific accumulation of GUS was maintained (Figure 5C). This finding implies that sub-region 2.1 contains a quantitative enhancer element. Deleting subregion 2.2 had no clear additional impact on either BS-specificity or activity (Figure 5A&D). However, the subsequent deletion of sub-regions 2.3 and 2.4 caused loss of GUS activity and also loss of GUS staining in BS and vascular tissue (Figure 5A,E&F). These data indicate that *cis*-regulatory elements mediating BS expression are situated in sub-region 2.3, or that quantitative elements in this region mask qualitative elements in distal sub-regions. To address these options, 3’ deletions of region 2 were also created (Figure 5G). As sub-regions 2.1 and 2.2 had little impact on BS expression, the last three sub-regions were fused to the minimal CaMV35S promoter. Expression from each of these three constructs was low (Figure 5G) however BS expression could be observed from the construct containing sub-regions 2.3, 2.4 and 2.5 (Figure 5H). Removal of sub-region 2.5 resulted in loss of GUS in rosette leaves although cotyledons still showed patchy staining restricted to the BS and vascular tissue (Figure 5I). Once sub-region 2.4 was removed GUS activity was further reduced and GUS staining was no longer detectable even in seedlings (Figure 5G&J). We conclude that sub-regions 2.3 and 2.4, which contain a total of 350 base pairs, contain *cis*-regulatory elements necessary for BS expression of *AtSULTR2;2*.

**Figure 5:**
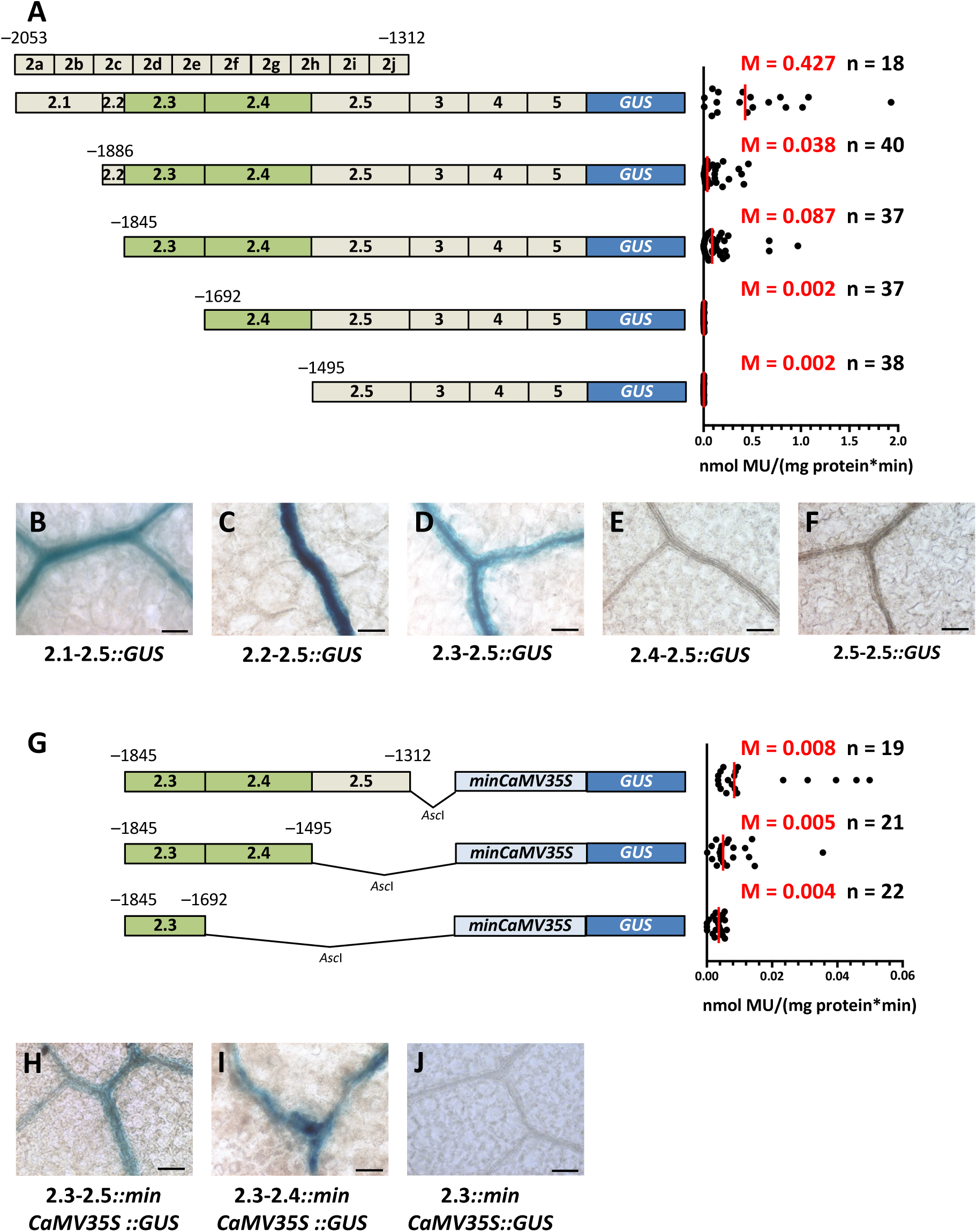
A short region of the *SULTR2;2* promoter that is necessary for expression in the bundle sheath. **(A)** Schematic of deletions made to region 2 of the *SULTR2;2* promoter (left) and quantitative analysis of expression from each construct based on the GUS activity assay (right). Activity was no longer detectable when subregions 4 and 5 were removed. **(B-F)** Histochemical staining of leaves indicated that subregion 3 is required for expression in the bundle sheath. **(G)** Schematic of 3’ deletion constructs placed upstream of the minimal CaMV35S promoter (left) and quantitative analysis of expression from each construct based on the GUS activity assay (right). **(H-**Nucleotides from −1845 to −1495 are sufficient to drive expression in the bundle sheath of leaves (H) and cotyledons (I) respectively. **(J)** Deletion of nucleotides −1692 to 1312 abolishes accumulation of GUS in the BS. Data from GUS activity assays include the median (M) indicated by red lines and the number (n) of independent lines. Histological GUS assays were allowed to proceed for 4 h (B), 47 h (C), 4 h (D) 6 d (E, F), 5 d (H), 2 d (I) and 29 h (J). Scale bars represent 50 µm.

The *cis*-regulatory elements necessary for BS-specific expression of *AtSULTR2;2* appear to be located in a 350 nucleotide region of the promoter. As finer-scale deletions had failed to identify the exact *cis*-elements responsible for this phenotype (Figure 4) a phylogenetic approach was undertaken. Orthologues of *AtSULTR2;2* were identified from seven species of the Brassicaceae. Alignments of sequences 5 kb upstream of each orthologue indicated that with the exception of *A. lyrata* that contains a 446 nucleotide insertion, region 2 is highly conserved (Figure 6). However, no short sequences or motifs within this sequence that may restrict expression to the BS could be identified (Figure 6). Although the results of this alignment therefore do not identify a specific *cis*-element that could be bound by a transcription factor responsible for generating BS expression, they do support the functional analysis and implicate the whole of region 2 as a critical component of the *SULTR2;2* promoter for BS expression.

**Figure 6:**
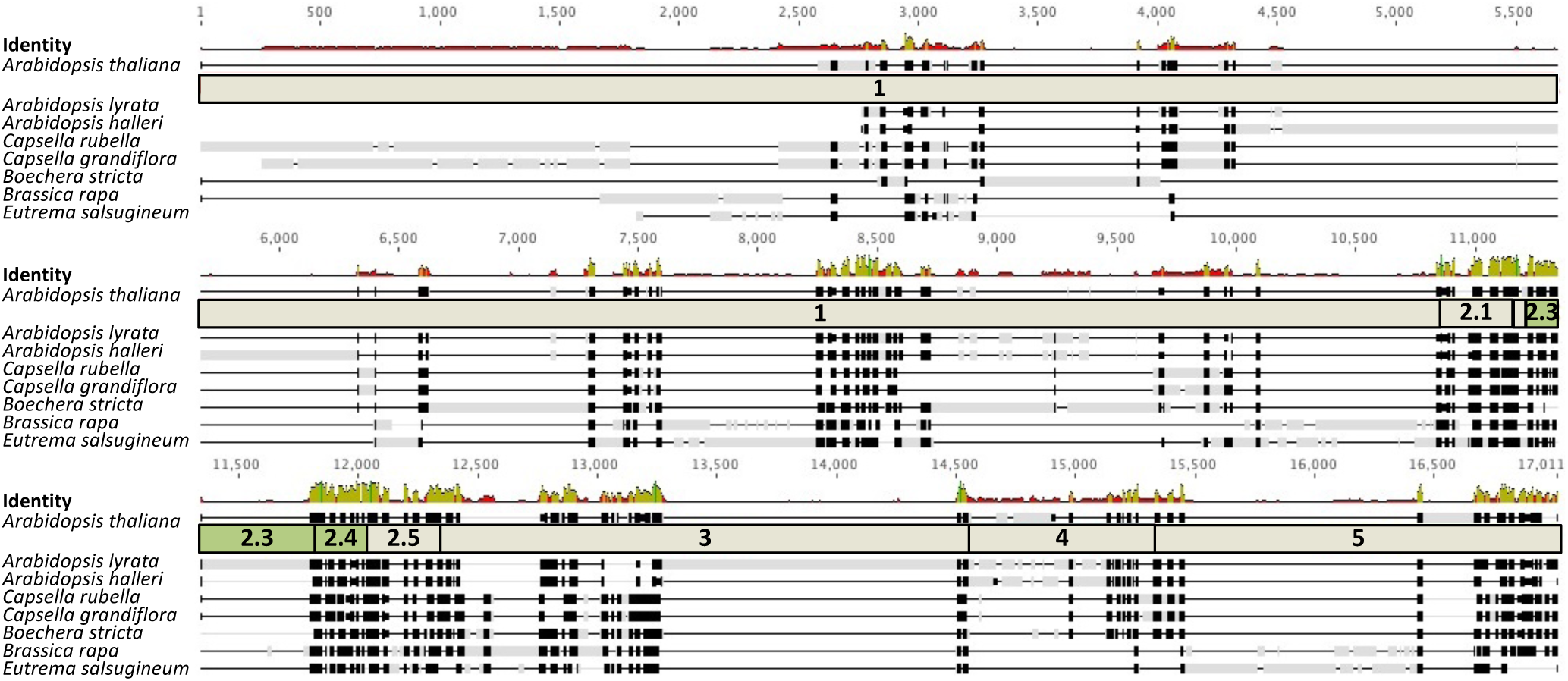
The region necessary for BS expression of *SULTR2;2* is highly conserved in the Brassicaceae. The promoter of *AtSULTRT2;2* was aligned against sequences ~5 kb upstream of genes from seven additional species of the *Brassicaceae*. With the exception of *Arabidopsis lyrata*, which contains a 446 bp insertion, region 2 is highly conserved. Black boxes indicate strong similarity of sequences, grey boxes sequences not matching the consensus sequence and black lines gaps. The level of similarity is also indicated on top of the alignment. High peaks in green mark strong similarity, low red peaks poor similarity.

### The *AtSULTR2;2* promoter is capable of driving BS specific expression in C_4_ *Flaveria bidentis*

As the *GLDPA* promoter from the C_4_ species *F. trinervia* is able to confer BS specific expression in C_3_ *A. thaliana* (Engelmann *et al*., 2008) we next tested whether the *A. thaliana SULTR2;2* promoter would lead to BS expression in C_4_ *F. bidentis*. GUS activity in transgenic *F. bidentis* plants was about fourfold higher than that in *A. thaliana* (Figure 7A&B). However, histochemical analysis of mature leaves revealed a very similar expression pattern to that in *A. thaliana* with strong GUS accumulation in BS and vascular tissue but not in mesophyll cells (Figure 7C). This indicates that transcription factors from *F. bidentis* recognise BS *cis*-regulatory elements from the Brassicaceae. The most parsimonious explanation for this finding is that these sequences represent part of an ancient and conserved mechanism that restricts gene expression to BS cells of dicotyledenous leaves.

**Figure 7:**
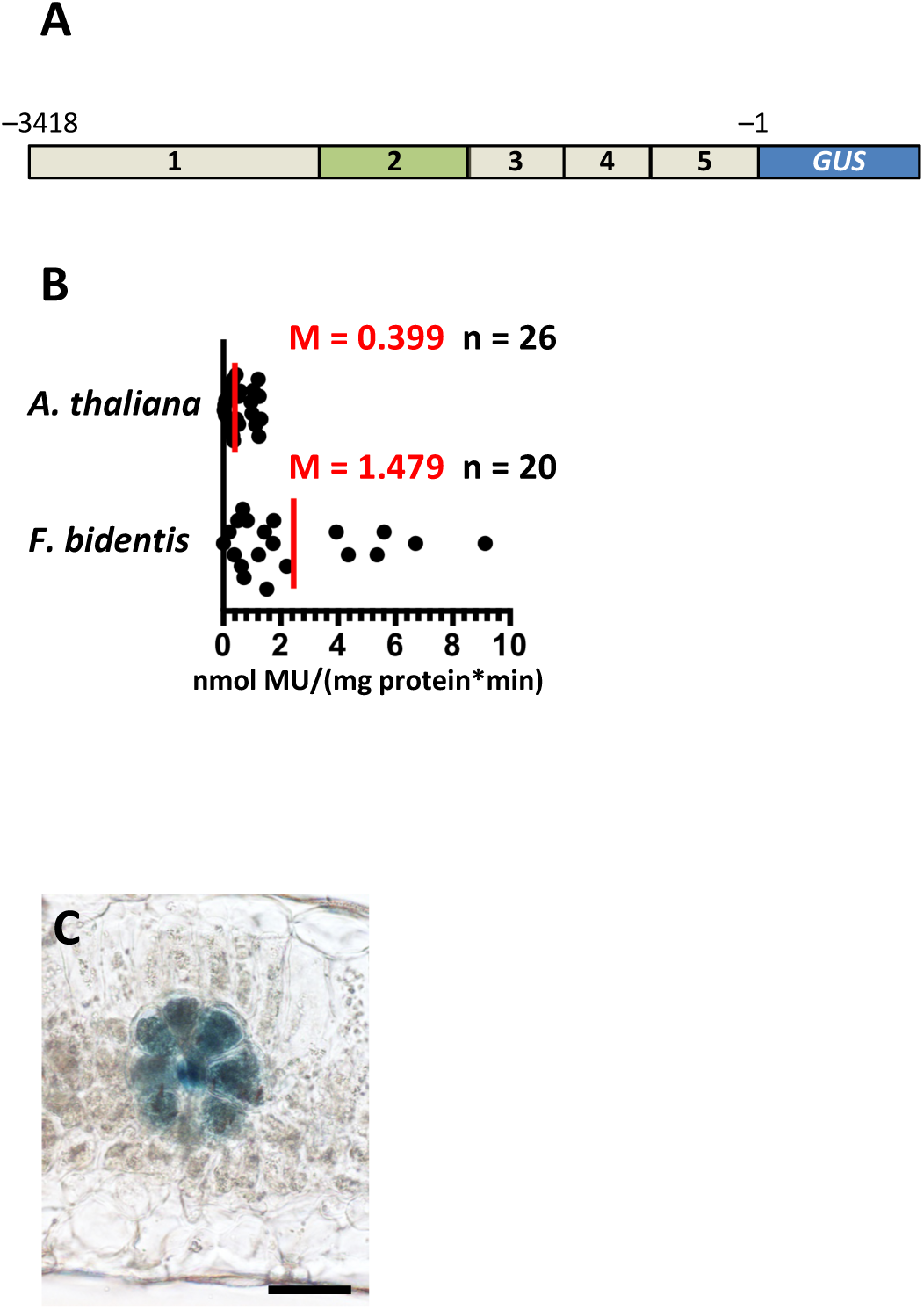
The *A. thaliana SULTR2;2* promoter generates strong bundle sheath expression in the C_3_ species *Flaveria bidentis*. **(A)** Schematic of the sequence placed into *Flaveria bidentis*. **(B)** Quantitative analysis of expression from each construct based on the GUS activity assay. To facilitate comparison GUS data from the same construct in C_3_ *A. thaliana* are included. **(C)** Representative image of transverse section of *Flaveria bidentis* after histochemical staining for GUS. Data from GUS assays include the median (M) indicated by red lines and the number (n) of independent lines. Histological GUS assays were allowed to proceed for 4 h. Scale bar represents 50 µm.

## Discussion

### *SULTR2;2* contains a short region that activates expression in the bundle sheath

*SULTR2;2* encodes a low affinity transporter that facilitates movement of sulphate from the vascular bundle to palisade cells in the leaf (Takahashi *et al*., 2000). Consistent with this function, analysis of both GUS and the histone2B::YFP fusion indicated that the *SULTR2;2* promoter directs expression to veins as well as to BS cells. It was notable that none of the various deletions we made to this promoter led to expression being restricted to either veins or bundle sheath cells. The *GLDPA* promoter from *F. trinervia* also drives expression in both veins and BS cells of *A. thaliana* (Engelmann *et al*., 2008). These findings therefore imply that the *SULTR2;2* and *GLDP* genes may be controlled by gene regulatory networks that are shared by these cell-types. It is possible that these cell-types share some gene regulatory networks because they are derived from the same lineage (Dengler and Nelson, 1999; Soros and Dengler, 2001).

Within the *SULTR2;2* promoter, one specific region that impacted on gene expression in the BS was identified. This sequence, consisting of 350 nucleotides, is both necessary and sufficient for restricting expression to this cell-type. Small consecutive deletions within this region failed to abolish this spatial patterning implying that multiple contiguous elements act redundantly to generate strong and stable expression in the BS. The only detectable impact of deleting any part of this region was for strength of expression to be reduced. We therefore propose that either multiple independent BS modules contained within this region act additively, or that distinct quantitative elements are co-located, and at least partially overlapping with, *cis*-elements that determine this cell-specificity. Redundancy of this sort has previously been reported for the promoter of *Phenylalanine Ammonia Lyase*2 that drives xylem-specific expression in tobacco (Leyva *et al*., 1992; Hatton *et al*., 1995), *DORNRÖSCHEN-LIKE* of *A. thaliana* that contains three functionally redundant enhancers (Comelli *et al*., 2016), and *EVEN-SKIPPED (EVE)* from *Drosophila melanogaster* where a minimal enhancer is sufficient to direct expression of *EVE* to the second stripe, but surrounding binding sites increase the robustness of this patterning during genetic and environmental perturbations (Ludwig *et al*., 2011). Thus, although the exact role of redundancy in the regulation of *SULTR2;2* in the BS is unclear, it may also increase robustness in the control of gene expression during environmental perturbations, and/or increase patterning precision (Barolo, 2012; Payne and Wagner, 2015).

Compared with other examples of elements that restrict gene expression to BS cells of C_3_ species, this single block of sequence from *SULTR2;2* that acts as a positive regulator of transcription appears to operate via relatively simple mechanisms. For example, the *F. trinervia GLDPA* generates BS-specific expression (Engelmann *et al*., 2008) because of a complex interplay between transcriptional and post-transcriptional processes. These are mediated by distal and proximal sequences relative to the translational start site, leading to repression of *GLDP* expression in mesophyll cells (Wiludda *et al*., 2012). We therefore propose that the positive regulator located upstream of *SULTR2;2* could be used as a synthetic module to manipulate or engineer processes in BS of *A. thaliana*.

### The bundle sheath element of *SULTR2;2* is conserved in the Brassicaceae and functional in the Asteraceae

Alignment of *SULTR2;2* promoters from multiple species of the Brassicaceae did not reveal an individual shared *cis*-element but rather highlighted sequence that was conserved across the whole of region 2. This sequence conservation argues for relatively strong purifying selection compared with the rest of the *SULTR2;2* promoter and also implies that this region may function as part of a widely conserved positive regulator of gene expression in the BS of the Brassicaceae. Consistent with this proposal and indicating that these regulatory elements may be even more ancient, when the *SULTR2;2* promoter was placed into the phylogenetically distant C_4_ species *F. bidentis* it was also recognized by *trans*-factors that restricted gene expression to the C_4_ BS. This is analogous to the behavior of the *GLDPA* promoter from C_4_ *F. trinervia* which is able to restrict expression to the BS of *A. thaliana* (Engelmann *et al*., 2008; Wiludda *et al*., 2012). Currently, the crown ages of the rosids and asterids are estimated to be 108-117 and 107-117 million years ago (Sanderson *et al*., 2004; Wikström *et al*., 2001) indicating that these clades diverged in the Early Cretaceous. Whilst for both *SULTR2;2* and *GLDP* it is possible that different mechanisms lead to BS specific expression in species of the Brassicaceae and Asteraceae it seems more likely that expression of each gene is determined by ancient and highly conserved *cis*-regulatory codes that that have been maintained since these clades diverged from their last common ancestor. Although the plant vasculature is thought to have started to evolve from 450 to 430 million years ago (Furuta *et al*., 2014; Ye, 2002) to our knowledge there are no clear estimates of when the BS originated. It would be intriguing if regulatory networks operating in both the veins and BS cells are uncovered that can be associated with the evolution of the vasculature in early diverging lineages of land plants.

## Materials and methods

### Cloning of promoter-reporter gene constructs and 5’ Rapid Amplification of cDNA ends

All DNA fragments created by PCR were confirmed by DNA sequencing. Generation of full-length promoter construct via PCR was constructed with *Arabidopsis thaliana* Columbia-0 (Col-0) genomic DNA. Subsequent constructs were generated using this as a template. Restriction sites were added to the respective fragments by PCR and fragments were inserted into pBI121 or a partially modified pBI121. Region 2 with internal deletions were synthesised by GenScript and swapped with the full-length region 2 of 2::5.2::*GUS* to generate the internal deletion constructs. Total RNA from leaves of wild type *A. thaliana* Col-0 plants was extracted, DNase I treated and purified with the RNeasy^®^ Plant Mini Kit (Qiagen, Hilden, Germany). cDNA was generated from 1 µg RNA and then 5’ RACE-PCR performed using the Advantage^®^ 2 DNA Polymerase Mix (Clontech Laboratories, Mountain View, USA) or Phusion^®^ HF DNA Polymerase (Thermo Fisher Scientific, Waltham, USA). Two nested 3’ oligonucleotides, AtSultr2;2-11 and AtSultr2;2-13, both binding in the cDNA of *AtSULTR2;2*, were used. PCR products were cloned and confirmed via colony PCR. Correct clones were subjected for plasmid preparation and sequencing.

### Transformation and plant growth

*A. thaliana* ecotype Col-0 was transformed using floral dipping (Clough and Bent, 1998; Logemann *et al*., 2006) using *Agrobacterium tumefaciens* strain AGL1. *F. bidentis* was transformed as described previously (Chitty *et al*., 1999). Successful transformations of *A. thaliana* or *F. bidentis* were tested by PCR. Before transplanting to soil, positive transformants of *A. thaliana* were selected on kanamycin. Seeds were sterilized by washing twice for 5 min with 20 % (v/v) DanKlorix (Colgate-Palmolive, New York City, USA) and 0.02 % (v.v) Triton-X 100 (Fluka Analyticals, Buchs, Switzerland) and four times with sterile water. Stratification of seeds was performed at 4 °C for 48 hours prior to spreading them on ½ MS pH 5.7 containing 10 g l^-1^ (w/v) sucrose (Sigma-Aldrich, St. Louis, USA), 0.5 g l^-1^ MES (Biomol, Hamburg, Germany), 2.13 g/l (w/v) Murashige-Skoog basal salts (Duchefa Biochemie, Haarlem, Netherlands), 0.75 % agar (SERVA Electrophoresis, Heidelberg, Germany), 50 µg ml^-1^ kanamycin (Sigma-Aldrich, St. Louis, USA) and 100 mg l^-1^ Cefotaxim (Fresenius Kabi Deutschland, Bad Homburg, Germany). Plants were transferred to 14 h light/10 h dark and temperatures of 23 °C day and 20 °C night and a light intensity of 90 µmol m^-2^ s^-1^.

### Visual and quantitative analysis of reporter genes

To take account of effects caused by transgene insertion into different genomic locations at between nine and forty-one independent T_0_ plants were analysed for each construct. Histochemical analysis was performed as described previously (Engelmann *et al*., 2008). For *A. thaliana* either three to four-week old rosette leaves or ten to fourteen-day old seedlings and for *F. bidentis* the sixth leaves of 40-50 cm tall plants were used. Transverse sections were prepared manually using a razor blade. Stained leaves were imaged using light microscopy. Quantification of GUS activity was performed via fluorimetric assay (Jefferson *et al*., 1987) using two to four leaves of three- to four-week old T_0_ *A. thaliana* plants or the fifth leave of 40-50 cm tall T_0_ *F. bidentis* plants, respectively. The Mann-Whitney U test was used to determine statistical differences between datasets. Imaging of H2B::YFP was performed on a Zeiss LSM 780 confocal laser-scanning microscope, and YFP fluorescence excited at 514 nm and emission detected between 517 and 569 nm.

## Acknowledgments

Work performed in PWs laboratory (SK, KP, MK, UG) was supported by the Deutsche Forschungsgemeinschaft through the Cluster of Excellence in Plant Sciences Duesseldorf-Cologne (EXC 1028). HW supported by a BBSRC PhD studentship. We thank the Center for Advanced imaging (HHU Düsseldorf) for provision and technical assistance of the Zeiss LSM 780 laser-scanning microscope.

## Figure legends

**Supplemental Figure 1: Additional deletion series generated for *SULTRT2;2*.(A)** Schematicsillustrating each deletion (left) and quantitative analysis of expression from each construct based on the GUS activity assay (right). The series was designed to remove around 400 bp each time and to avoid *cis*-regulatory elements predicted by the software PLACE (Higo *et al*., 1999). Deletion of region II resulted in a decline in GUSactivity and deletion of region III lead to a total loss of expression. **(B-H)** Representative images after histological staining for GUS indicating that deleting region III led to loss of GUS in the bundle sheath. Data from GUS activity assays include the median (M) indicated by red lines and the number (n) of independent lines. Histological GUS assays were allowed to proceed for 6 h. Scale bars each 50 µm.

**Supplemental Figure 2:** Alignment of 5’ ends of cDNAs obtained via 5’ Rapid amplification of cDNA ends. No distinct transcription start site was found. The top line shows the template used for alignment. Predicted translational start sites are marked in blue (annotated in TAIR Accession 1009028759) and red (from Takahashi *et al*. (2000)). A TATA box motif found by the PLACE database (Higo *et al*., 1999) is underlined. The black arrowhead marks the TAIR annotated transcription start site (TAIR Accession 1009028759).

